# MirGeneDB 3.0: Improved taxonomic sampling, uniform nomenclature of novel conserved microRNA families, and updated covariance models

**DOI:** 10.1101/2024.09.27.615356

**Authors:** Alexander W. Clarke, Eirik Høye, Anju Angelina Hembrom, Vanessa Molin Paynter, Jakob Vinther, Łukasz Wyrożemski, Inna Biryukova, Alessandro Formaggioni, Vladimir Ovchinnikov, Holger Herlyn, Alexandra Pierce, Charles Wu, Morteza Aslanzadeh, Jeanne Cheneby, Pedro Martinez, Marc R. Friedländer, Eivind Hovig, Michael Hackenberg, Sinan Uğur Umu, Morten Johansen, Kevin J. Peterson, Bastian Fromm

**Author notes:** contributed equally (Alexander W. Clarke, Eirik Høye). corresponding authors &. University of Georgia, Athens, GA, USA.

## Abstract

We present a major update of MirGeneDB (3.0), the manually curated animal microRNA gene database. Beyond moving to a new server and the creation of a computational mirror, we have expanded the database with the addition of 33 invertebrate species, including representatives of 5 previously unsampled phyla, and 6 mammal species. MirGeneDB now contains entries for 21, 822 microRNA genes (5, 160 of these from the new species) belonging to 1743 microRNA families. The inclusion of these new species allowed us to refine both the evolutionary node of appearance of a number of microRNA genes/families, as well as MirGeneDB’s phylogenetically informed nomenclature system. Updated covariance models of all microRNA families, along with all smallRNA read data are now downloadable. These enhanced annotations will allow researchers to analyze microRNA properties such as secondary structure and features of their biogenesis within a robust phylogenetic context and without the database plagued with numerous false positives and false negatives. In light of these improvements, MirGeneDB 3.0 will assume the responsibility for naming conserved novel metazoan microRNAs. MirGeneDB is part of RNAcentral and Elixir Norway, and is publicly and freely available at master.cloud.mirgenedb.org.

**Key Points:**

1. Major update to the manually curated and uniformly named microRNA gene database MirGeneDB
2. 114 animal species, >1700 microRNA families and ∼20 000 genes searchable, browsable and downloadable
3. New features to download all smallRNA read data and updated covariance models for each family

## Introduction

The last five years have seen the publication of more than 55, 000 papers referencing microRNAs in their title or abstract, more than any other type of non-coding RNA. These studies, however, continue to be challenged by long-standing problems that diminish the accuracy of publicly available small RNA annotations (1–3). Previous studies have found that as many as two-thirds of the purported microRNAs recorded in public databases are likely false positives (4–18), and that existing nomenclatural schemes often failed to reflect the real evolutionary histories of a number of important microRNA families. These errors hinder comparative work and have led to a confusing proliferation of study-specific databases (5, 19–24). Further, they limit the power of microRNA data and the replicability of studies that use them across all branches of the biological sciences, from taxonomy and phylogenetics (25) to biomedical research (26).

Fortunately, these issues arise, not from an inherent problem in the study of microRNAs *per se*, but from the fact that many databases simply catalog the microRNAs described in the published literature without independently curating them. Although the process of reviewing purported microRNAs by hand is labor-intensive and cannot guarantee the absence of false negatives, especially in the case of tissue-specific or poorly expressed genes, it can eliminate virtually all false-positive results, significantly improving the reliability of microRNA data. To this end, in 2015, we established MirGeneDB, the first publicly accessible, manually curated microRNA gene database (13). Expanding on the pre-NGS criteria for microRNA annotation (27), we established a rigorous and consistent set of criteria to annotate a high-confidence set of microRNAs across the metazoa. Over the last nine years, MirGeneDB’s repertoire has grown from just 4 (13), to 45 (18), to 75 species (17), including model and non-model systems drawn from roughly two-thirds of all animal phyla. Recently, we (28) and others (29) demonstrated that this well-curated data can be used to train algorithms to predict conserved microRNAs from genomes only, highlighting MirGeneDB’s value and potential for both comparative genomics and phylogenetics (25).

Despite its regular expansions, major branches of the metazoan tree were either wholly unrepresented or undersampled in previous versions of MirGeneDB, and, as a result, we have not been able to fully capture the patterns of microRNA evolution within several clades. To improve MirGeneDB’s cross-section of the metazoa and address residual nomenclatural and modeling issues, we have completed a third major update, MirGeneDB 3.0. This update enhances MirGeneDB’s comparative data set with more than 5000 new genes and more than 200 new gene families from 39 new species, drawn largely from invertebrate clades. Altogether, more than 20, 000 accurately annotated, consistently named and curated microRNA genes from 114 metazoan species can now be browsed, searched and compared in MirGeneDB. We show that this data is useful for identifying the structural and sequence features of metazoan microRNAs, with some noteworthy outliers, with implications for microRNA prediction and annotation. We further provide downloadable covariance models (CM) and processed read files for all species. To address the present lack of an institution that names novel microRNAs, and to provide a functional reference, MirGeneDB will now assume the responsibility of naming novel microRNAs that are conserved between at least two animal species. It is our hope that this expansion and our efforts to name novel genes and families will allow MirGeneDB to have an even wider user base and continue to stand as the “gold standard” (30–37) for metazoan microRNA annotation across the biological sciences.

### Improving Phylogenetic Representation across the Metazoa

The new species included in this update extend MirGeneDB’s sample of the metazoans and refine its resolution within several clades. These improvements are most noticeable among the invertebrates, which account for 33 of the 39 new taxa (Figure 1, blue species). At the broadest scale, we have added members of five previously absent phyla: Acoela (represented by *Hofstenia miamia* and *Symsagittifera roscoffensis*), Nematomorpha (*Gordionus* sp.), Nemertea (*Lineus longissimus*), Phoronida (*Phoronis australis*), and Priapulida (*Priapulus caudatus*). More narrowly, we have expanded our existing sample of several phyla. We added four new annelid species: the tubeworm *Owenia fusiformis*, the progenic orbiniid *Dimorphilus gyrociliatus*, the ragworm *Platynereis dumerilii*, and the peanut worm *Sipunculus nudus*. Similarly, we added eight new species of molluscs: the solenogasters *Epimenia babai* and *Wirenia argentea*, the monoplacophoran *Laevipilina hyalina*, the polyplacophorans *Mopalia mucosa* and *Acanthopleura granulata*, and the scaphopod *Pictodentalium vernedei*, along with second representatives for the gastropods (*Haliotis rufescens*) and the bivalves (*Ruditapes philippinarum*). We also expanded our representation of the platyhelminths with the free-living polyclad *Prostheceraeus crozieri* and three parasitic neodermatans: the monogenean *Gyrodactylus salaris*, the trematode *Schistosoma mansoni*, and the cestode *Echinoccocus granulosus*. To the syndermatans (rotifers), we added the monogonont *Brachionus koreanus* and the bdelloid *Adineta vaga* (both free-living), as well as the ectoparasitic seisonid *Seison nebaliae* and the endoparasitic acanthocephalans *Neoechinorhyncus agilis* and *Pomphorhyncus laevis*. We also annotated four parasitic arthropods: the mosquito *Anopheles gambiae*, the tsetse fly *Glossina pallidipes,* the parasitic wasp *Diachasmimorpha longicaudata*, and the spider mite *Tetranychus urticae*. Finally, to better understand the evolution of microRNAs in cnidarians, we added two additional anthozoans: the stony coral *Acropora digitifera* and the sea anemone *Actinerus* sp.

**Figure 1:**
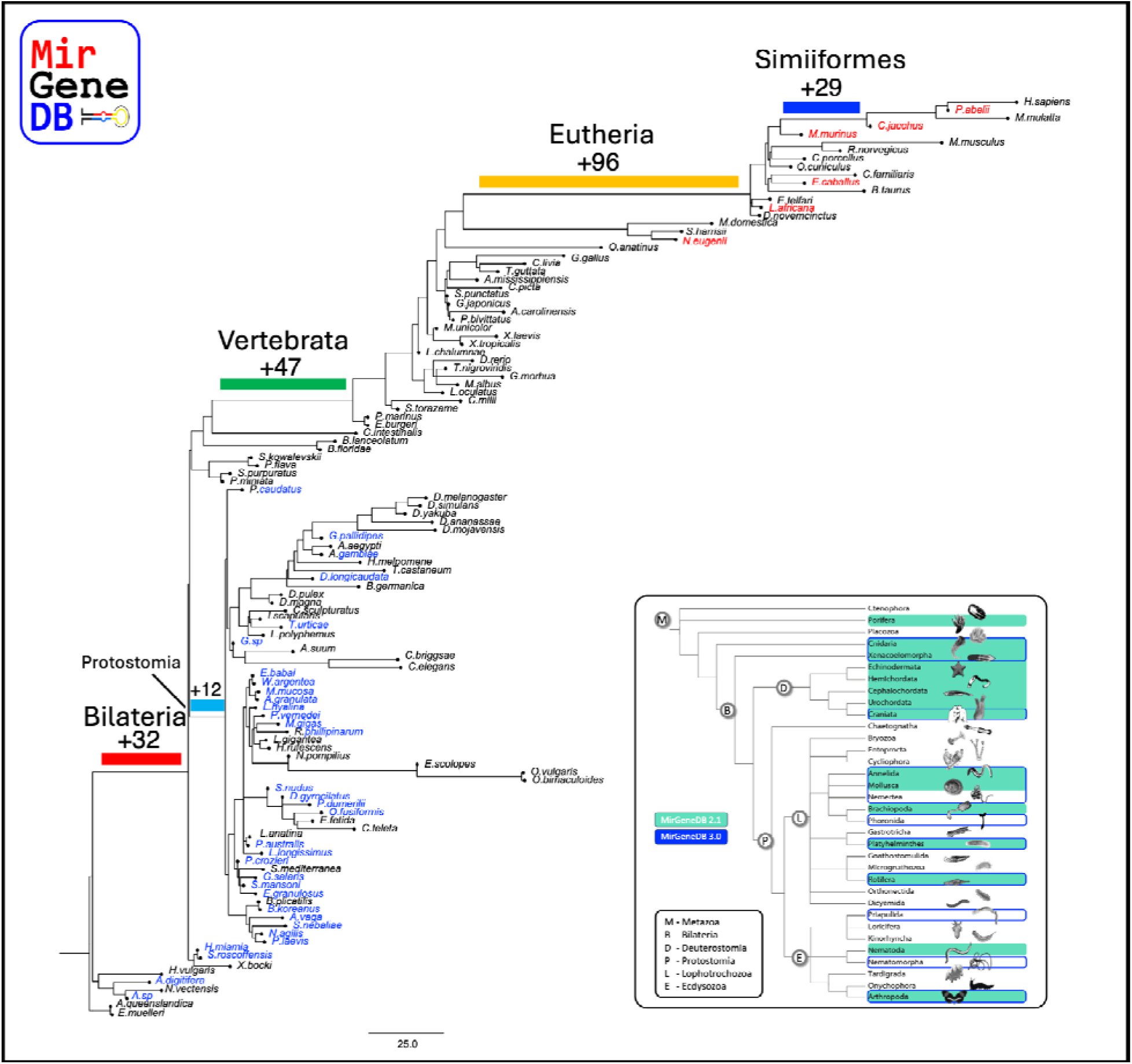
The evolution of 1743 microRNA families across the 114 metazoan species annotated in MirGeneDB 3.0, with branch lengths corresponding to the total number of microRNA family-level gains plus family-level losses. New species added in this release are in blue (invertebrates) and red (vertebrates). Colored bars indicate some of the phylogenetic nodes associated with bursts of family-level microRNA innovation, with the number of new families at that node below. Inset: simplified metazoan phylogeny to highlight differences in MirGeneDB representation at the phylum level.

In addition to their phylogenetic significance, these new taxa enhance MirGeneDB’s coverage of several ecologies. For example, the secondarily miniaturized *Dimorphilus* and the three non-acanthocephalan rotifers double MirGeneDB’s suite of meiofaunal species (formerly consisting of only the rotifer *Brachionus plicatilis* and the two *Caenorhabditis* species). Among these taxa, *Dimorphilus* and *Seison* are especially interesting because they have the shortest genomes in our database—at only 70 Mbp (30) and 43 Mbp (38), respectively. This update also expands MirGeneDB’s repertoire of parasites with new micropredators (*Anopheles* and *Glossina*), trophically transmitted parasites (*Schistosoma*, *Echinococcus*, and the acanthocephalans), and directly transmitted parasites (*Gyrodactylus*, *Tetranychus*, and *Seison*). With the two parasitoids (*Diachasmimorpha* and *Gordionus*) in this update, MirGeneDB now contains representatives for 4 of the 6 canonical parasitism strategies (39).

Along with these invertebrates, MirGeneDB 3.0 now contains 6 additional mammalian species (Figure 1, red species). Previous studies have found that the rate of family-level microRNA innovation increased in the primates, relative to other mammals (29). To more precisely resolve the timing of this burst, this update includes 3 new species: *Pongo abelii*, *Callithrix jacchus*, and *Microcebus murinus*, representing a non-human hominoid, a platyrrhine, and a strepsirrhine, respectively. Together, these species indicate that the primate innovation pulse began within the Simiiformes and continued into the Haplorhini (Figure 1, dark blue bar). This update will also introduce the first members of Perissodactyla (*Equus caballus*), Paenungulata (*Loxodonta africana*), and Diprotodontia (*Notomacropus eugenii*) to MirGeneDB.

Overall, the new species included in MirGeneDB 3.0 increase its comparative power and improve our ability to resolve the deep-time evolutionary dynamics of microRNAs families and genes (40, 41). Indeed, by including these new species, we were able to reevaluate the evolutionary origins of several microRNA families and establish the conservation (and biological importance) of many lineage-specific families, especially among the Annelida, Mollusca, Syndermata, and Platyhelminthes. Altogether, the 720 microRNA families now known to be conserved between two or more species support the monophyly of 80 different higher-level taxa (Supplementary Table 1).

Furthermore, the continual inclusion of new species into MirGeneDB buttresses our original insights (42–44) that new microRNA families continually appear in most metazoan lineages, with gains outweighing losses in most taxa outside of a few parasitic and/or meiofaunal species. Indeed, these important patterns—found only with microRNAs and not with any other type of known regulatory molecules (41)—are robustly illustrated using banner-plots (Figure 2). By highlighting important phylogenetic nodes, the conservation of the number of microRNA families and paralogues across metazoans, as well as the potential usefulness of microRNAs as taxonomic and phylogenetic markers, is obvious.

**Figure 2:**
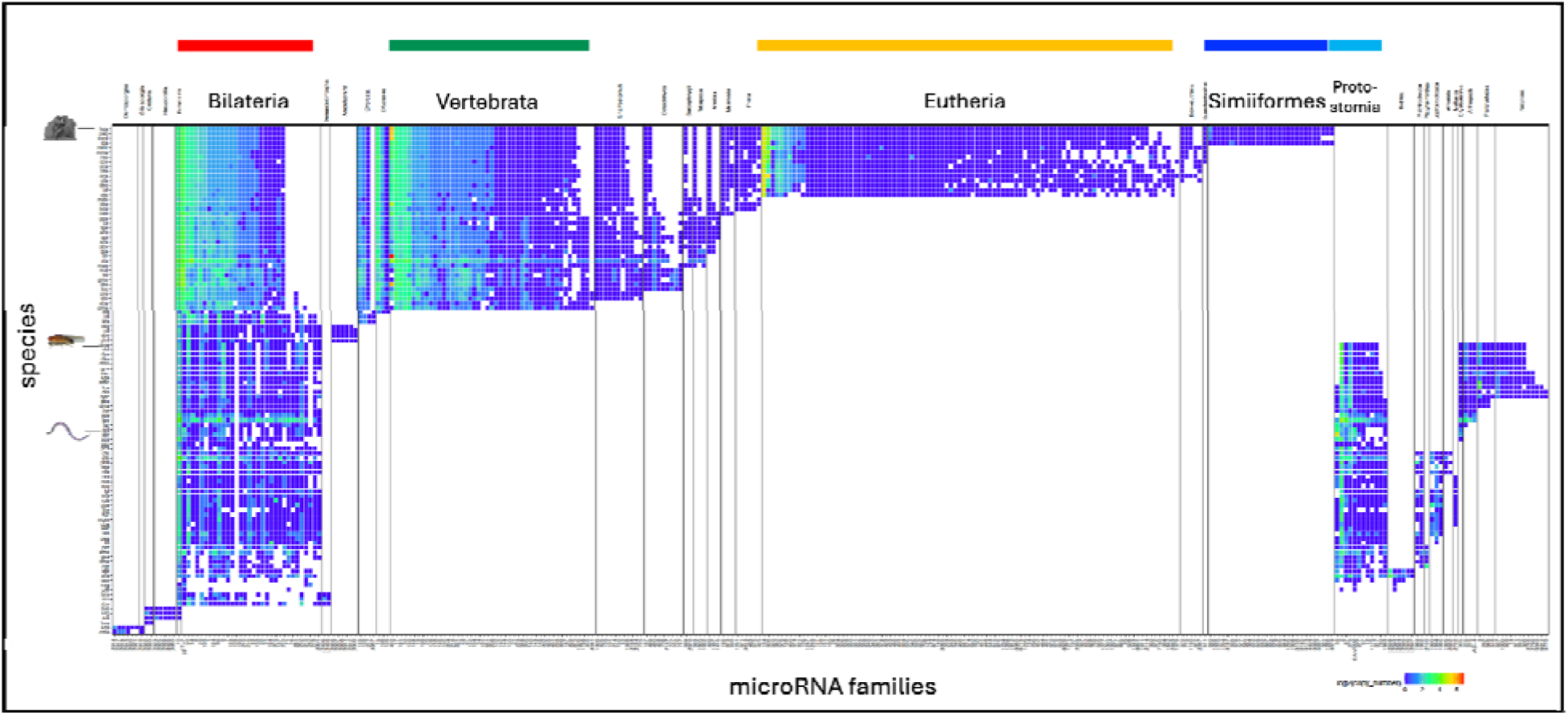
Banner plot of the 114 Metazoan microRNA complements in MirGeneDB 3.0. Rows represent species, sorted as in Figure 1, and columns represent microRNA families, sorted by node of origin and overall paralogue number (heatmap function, with blue indicating few paralogues and red/yellow indicating many). The coloured bars along the top depict the selected phylogenetic nodes highlighted in (Figure 1).

### Revising MicroRNA Cluster Annotation

Starting with this update to MirGeneDB, we will institute a new procedure for naming the members of microRNA gene clusters. Because it relied on miRBase, which was not informed by evolutionary history, our original system for annotating paralogous microRNA genes could not be effectively applied to large gene clusters. One example, among many, of the problems facing this system can be found in the important eutherian-specific microRNAs of the imprinted MIR-154/MIR-376 cluster on human chromosome 14 (45). In humans, this locus, which sits just downstream of three paternally expressed protein-coding genes, and 99 maternally-expressed small RNAs (including 51 microRNAs derived from lncRNAs, Figure 3), contains 33 MIR-154 genes, five MIR-376 genes, and two genes that constitute their own families (MIR-134 and MIR-541). In previous iterations of MirGeneDB, miRBase’s original Mir-154 gene was named Mir-154-P1, and subsequent genes in that family received paralogue IDs according to the order of their miRBase names (e.g., miRBase’s Mir-299 and Mir-323 received the MirGeneDB names Mir-154-P2 and Mir-154-P3). Though this system was relatively simple, it meant that a brief review of the cluster would give no indication of which paralogues might be novel in a particular species and which ancestral paralogues might have been lost.

**Figure 3:**
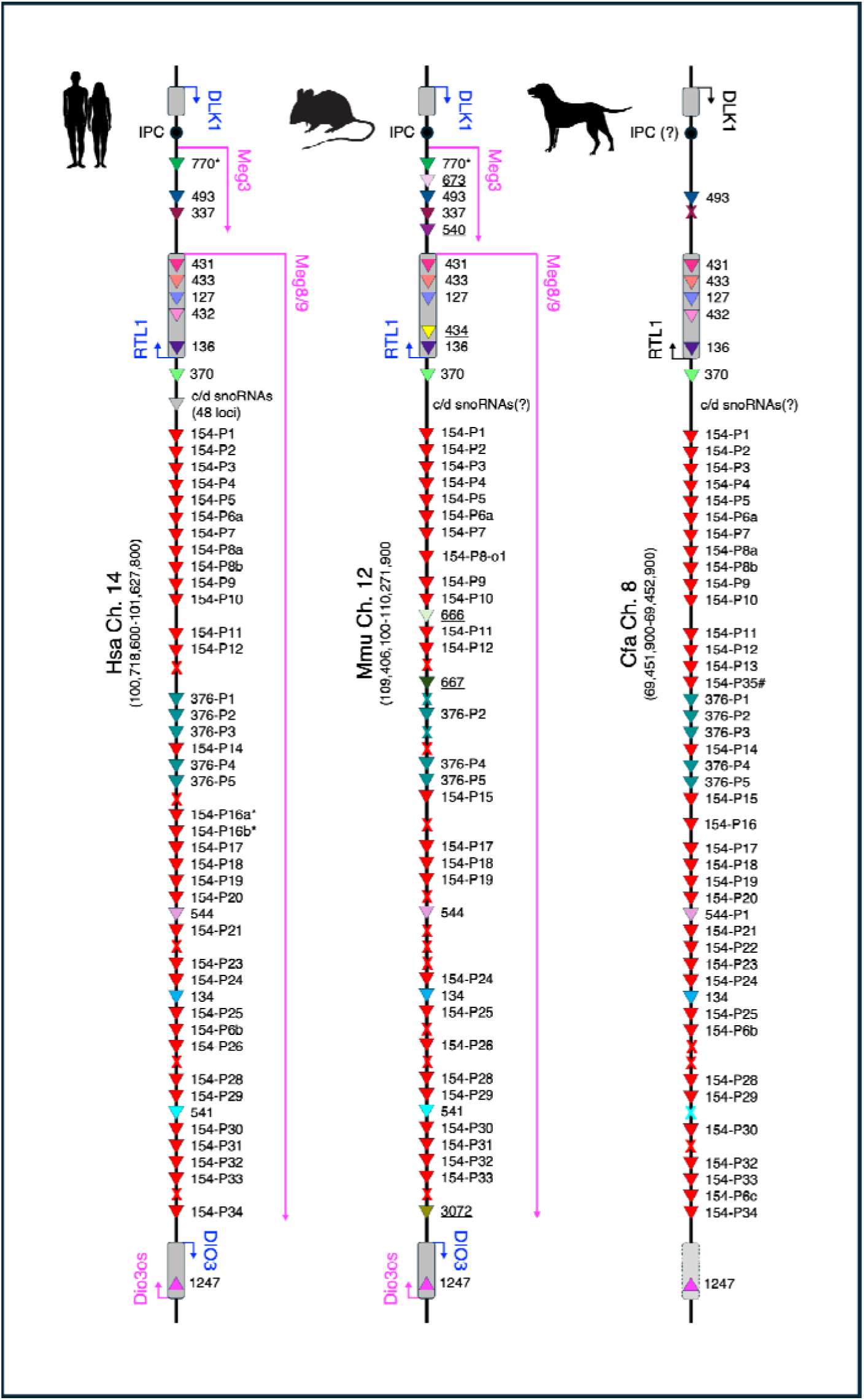
The annotation and nomenclature system for clustered microRNA loci used in MirGeneDB 3. Exemplified in the imprinted MIR-154/MIR-376 cluster on human chromosome 14 and in comparison to mouse and dog the 5’ -3’ naming scheme is shown.

To address this issue, MirGeneDB 3.0 will institute a 5’ to 3’ system for paralogue annotation, with the 5’-most gene in a cluster receiving the “P1” designation (for the MIR-154 family, this is miRBase’s mir-379, our old Mir-154-P7), with humans as the reference taxon. This system, when applied consistently, can immediately reveal both a species’ novel gene copies through additions to the paralogue number (e.g., Mir-154-P6a and P6b in humans) and its secondary losses of ancestral copies through absences of expected paralogue numbers (e.g., both Mir-P21 and -P23 are present in human, but -P22 is not, indicating that this gene must be present in other eutherians like the dog, Figure 3, far right). When this cluster is compared across multiple eutherian species, the loss of Mir-154 genes in murid rodents, for example, is striking (Figure 3, middle). Moreover, unique genes present in specific eutherian lineages that evolved after the eutherian LCA are now readily apparent by the fact that they have a higher paralogue number than their neighbors. For example, Mir-154-P35 sits immediately 3’ of Mir-154-P13 and 5’ of the Mir-154-P14 gene in dogs. Because humans are the reference taxon for the nomenclature, this indicates to the user that, instead of this gene being lost in humans, in which case the paralogue numbers would follow a normal ordinal sequence, it must have evolved somewhere in the lineage leading to dogs (in fact, it is present in the three laurasiatherians: *Canis*, *Bos*, and *Equus*). We have applied this system to all known microRNA clusters, with the exception of a few ancestral clusters like the LET-7 and MIR-96 clusters, making the annotation and the evolutionary understanding of these large clusters much easier.

### Nomenclatural Responsibilities

Historically, MirGeneDB has deferred to miRBase in certain aspects of nomenclature. Though we have always categorized microRNAs according to shared evolutionary history, rather than shared seed sequences (leading, for example, to the inclusion of miRBase’s mir-100 and mir-125 into the MirGeneDB MIR-10 family), we have previously avoided naming entirely novel microRNA genes or families ourselves. Instead, newly discovered sequences have generally received a provisional name containing their clade of origin, the word “novel,” and an ordinal ID (e.g., or CAT-Novel-1 for the first novel found in the Catarrhini) until at least one member of that family was added to miRBase and given a name there. Beginning with this update, however, we will depart from this policy.

At the time of writing, miRBase has not received a major update in 5 years (46), despite the recent release of a great number of small RNA datasets from dozens of previously unstudied species. In this time, MirGeneDB has been updated thrice (including this update) and, as a result, contains annotations for 52 metazoan species and several major clades, such as the cephalopods, afrotheres, and rotifers, that are not included in miRBase. With the enhanced phylogenetic resolution afforded by the 3.0 update, we have also been able to conduct a systematic review of gene-level relationships within several invertebrate microRNA families, refining our previous assessments of paralogy. For these reasons, MirGeneDB will, henceforth, assume primary responsibility for naming new microRNA families and genes in metazoans.

For a microRNA family to receive an official name in MirGeneDB, it will need to meet the rigorous biogenesis criteria described in earlier publications (13, 17, 18) and must be conserved between two or more species. Those novel families that appear to be unique to a single species will receive a provisional name in the format described above, with a three-letter species code as their clade-of-origin (e.g., Cte-Novel-1 for the first novel found uniquely in *Capitella teleta*), until their conservation is demonstrated through the sampling of additional species. The only exception to this general rule will be those species-specific microRNA families that have already received names from miRBase (e.g., Cel-Mir-792); such families will retain their current names.

To prevent discordances between previous and future microRNA nomenclature, MirGeneDB will continue to include miRBase names, when they exist, alongside our own nomenclature for microRNA genes and families. Furthermore, it will still be possible to search for miRBase IDs along with MirGeneDB IDs on our web server.

In light of our new responsibilities, we will also amend some of MirGeneDB’s internal policies. Prior to this release, gene names in MirGeneDB have been subject to continuous revision, both to address our own errors, as revealed by improved taxon sampling, and to facilitate the analysis of large microRNAs clusters. In the long run, however, the instability that these revisions create would make it difficult to use MirGeneDB effectively. For this reason, MirGeneDB will institute a naming freeze on all known microRNA genes and families. Going forwards, we will not rename genes in MirGeneDB, nor will we change the names of valid families that have received a name from miRBase. If future expansions of this database reveal that we have misunderstood the evolutionary histories of any known genes, we will make a note in the “Comments” section of the gene’s page, but will not otherwise alter its ID. In addition, we will not reuse the miRBase name of a rejected family for any valid one discovered in the future (e.g., the names MIR-68, MIR-69, and MIR-198 will never be used). The primary exception to these rules are so-called orphan genes, which are members of a microRNA family whose paralogy is fundamentally uncertain at this time. These genes, designated with a “–o,” will be renamed as the inclusion of new taxa refines our understanding of each major clade’s ancestral microRNA complement.

### Covariance Models

MirGeneDB’s rigorous curation criteria significantly improves the quality of microRNA annotation but requires significant human and computational resources. Furthermore, they prevent us from assessing the microRNA complements of species with public genomes but no public small RNA sequencing data. To overcome these limitations, we developed MirMachine in 2023 (28). The first version of MirMachine is based on MirGeneDB 2.1 and uses covariance models of 508 conserved microRNA families to predict which of those families are present in a given genome. In effect, this program enables us to qualify a species’ repertoire of conserved microRNAs, even when no small RNA data are available. These predictions can also be used to uncover microRNA pseudogenes, determine the conserved complements of extinct species, assess the quality of a genome assembly, and improve the performance of smallRNA-based microRNA prediction software. On average, these predictions have an accuracy of 97.5%, making MirMachine the first tool that can accurately predict microRNAs from metazoan genomes only (see also (29) for a human microRNA families focussed tool).

Despite MirMachine’s high overall accuracy, we recognized a potential to significantly improve its performance. Most importantly, the covariance models underlying the program tend to favor the patterns of variation and conservation observed in well-sampled clades, relative to more sparsely sampled ones. As a result, the models are more likely to make Type II errors (i.e., spuriously fail to predict an existing gene) when faced with otherwise unimportant sequence or structural changes in groups that were poorly represented, or unrepresented entirely, in MirGeneDB 2.1.

To resolve these issues, we have retrained the MirMachine covariance models using MirGeneDB 3.0’s expanded complement of species. Most importantly, the many new protostomes included in this update will diminish the influence that any single species or clade in this group has on the performance of the covariance models, reducing their false-negative rate. Similarly, the thousands of new genes in the training set have increased the total sequence diversity for all microRNA families, which will make the covariance models more tolerant of any new variation which they may encounter in a future genome. The new covariance models are publicly available on MirGeneDB 3.0 and will be included in a later, fully overhauled release of MirMachine.

### Sequence and conservation propensities of microRNAs

Though they are a powerful tool, curated datasets are sometimes too small to reliably capture the general characteristics of complex biological entities and their evolutionary histories. With this in mind, we have made a concerted effort to rigorously and exhaustively curate the (conserved) microRNA complements of as broad a set of metazoans as possible. As of this update, MirGeneDB covers the majority of phyla in the metazoan tree of life, often with several representative species, and samples a wide range of lifestyles, ecologies, and genomic characteristics, including dramatic differences in genome size and content (Peterson et al. 2024), and various numbers of whole genome duplication events (Peterson et al. 2022). As a result, we now have the ability to systematically examine the characteristics that unite microRNAs across the metazoa.

At the most basic level, we can provide robust length criteria for the precursor microRNA sequence as a whole, as well as the lengths of the processed arms. Over the last several decades, authors have used a bewildering array of ranges to describe these sequences. For example, a brief review of the published estimates for the range of processed arm lengths returns at least 10 different possibilities: 18-22, 18-23, 19-22, 19-23, 19-24, 19-25, 20-22, 20-23, 20-24, and 20-25 nt. The data in MirGeneDB allow us to conclusively state that the average lengths of both the 5p and 3p arms are 22 ± 1 nt. Importantly, no known, robustly curated microRNA has an arm length of less than 20 nt or more than 27 nt (and the few known arm products that are 27 nt long all belong to vertebrate-specific paralogues of the MIR-34 family). This characteristic should be used to help discriminate potential novel microRNA sequences from the many other types of non-coding RNA molecules present in animal cells. Because the mean length of the inter-arm loop is 16 nt, the mean length of a metazoan pre-microRNA sequence is 60 nt. The lengths of canonical precursor sequences, however, vary widely between clades, with the cnidarians having some precursors as short as 49 nt and the arthropods having some as long as 250 nt (Supplementary Figure 1). Indeed, the level of taxonomic inclusion in MirGeneDB 3.0 makes it obvious that some species have tightly constrained pre-microRNA lengths of less than 80 nt (e.g., vertebrates, nematodes, mollusks) while others appear to have no upper limit to the length to the loop (e.g., arthropods, platyhelminthes, rotifers). Although Fromm et al. (13) speculated that there might be a relationship between loop lengths and the number of Dicer genes in a species’ genome such that only taxa with 2 or more Dicer genes had unconstrained loop lengths, it is now known that even some taxa with only a single Dicer gene (e.g. Annelida, (47)) occasionally have extraordinarily long pre-microRNA sequences (e.g., Mir-1994 in the oligochaete annelid E. *fetida* that measures 137 nt in length). For this reason, the cause of the observed constraint in pre-microRNA length in vertebrates and other groups must remain an outstanding and enigmatic question.

**Table 1.**

|  | 5p arm |  |  |  | loop |  |  |  | 3p arm |  |  |  | base-pairing |
| --- | --- | --- | --- | --- | --- | --- | --- | --- | --- | --- | --- | --- | --- |
|  | min | median | max | SD | min | median | max | SD | min | median | max | SD |  |
| 5p Matures<br>(N=9'497) | 20 | 22 | 27 | 0.89 | 5 | 16 | 235 | 7.58 | 20 | 22 | 26 | 0.72 | 19±1.587 |
| 3p Matures<br>(N=10'940) | 20 | 22 | 27 | 0.99 | 7 | 15 | 238 | 8.27 | 20 | 22 | 26 | 0.81 | 19±1.581 |
| co-Matures<br>(N=410) | 20 | 23 | 26 | 0.96 | 6 | 16 | 170 | 5.92 | 20 | 22 | 26 | 0.77 | 19±1.647 |

Beyond the lengths of pre- and processed microRNA arm sequences, we can also now discern both sequence as well as structural propensities in metazoan microRNA sequences, in addition to processing propensities whereby specific trends might be found with either 5p-derived mature sequences versus 3p-derived mature sequences. Figure 4 shows heat maps for both from representative species, with sequence propensities in color and structural propensities in gray (See Supplementary Figure 2 for all species). These trends are determined from an analysis of all 20847 microRNA sequences in MirGeneDB 3.0.

**Figure 4:**
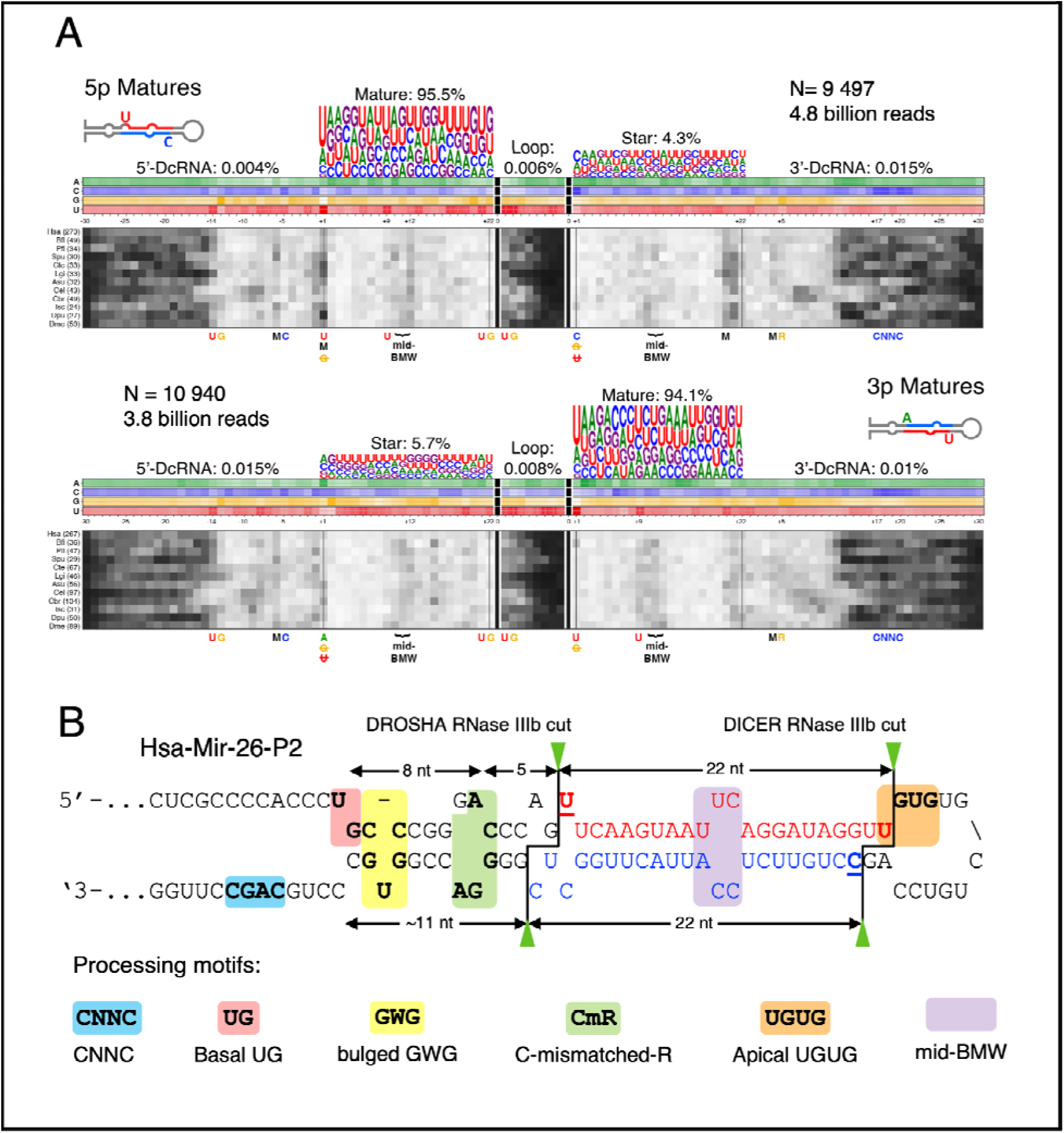
Global summary of the sequence and structure propensities of metazoan microRNAs. A. Sequence and structural signatures for 9497 5p mature microRNAs, and at the bottom for 10940 3p mature microRNAs for 114 bilaterian taxa (the number of considered microRNAs for the selected species is given in parentheses). Shown across the top are the read percentages from MirGeneDB split into matures, stars, loops, as well as 5’ and 3’ Drosha-cleavage (DcRNA) RNAs. Below the sequence logos are sequence-signature heatmaps (A = green; C = blue; G = orange; U = red); the darker the shade the higher the incidence of that particular nucleotide at that particular position. Below the sequence-signature heatmaps are the structural-signature heatmaps – the darker the shade the more likely the structure is unpaired at that position. Loop sequences are truncated to the minimal 8 nucleotides in required length (Fromm et al. 2015) for all sequences and for all taxa, and only show the first four 5’ nucleotides and the terminal 3’ nucleotides of each loop sequence. Note that both 5p mature and 3p matures have a high propensity to have a uridine in positions one and nine (see also the corresponding sequence logos). Interestingly, the stars belonging to 5p matures have a propensity to start with a cytosine at position 1, whereas stars belonging to 3p matures tend to start with an adenosine. Of note are the conspicuous unpaired regions immediately 5’ on the 5p arm and 3’ on the 3’ arm of the microRNAs found in *C. elegans* and *C. briggsae*, especially the ones whose mature gene products arise from the 5p arm, as well as the internal base-paired regions, in relation to all other analyzed bilaterian species. B. The sequence and structural propensities and processing motifs of a representative microRNA (Hsa-Mir-26-P2). Six processing motifs are clearly seen in many pri-microRNAs (moving 5’-3’ along the primary RNA sequence) and most are readily apparent with the sequence and structural propensities shown in A: 1) the basal or proximal UG motif (red) at the transition from the flanking unpaired sequence to base-paired stem structure; 2) the “bulged GWG” motif (yellow) with a bulged uridine or adenosine (W) on the 3p arm flanked by 5’C-G-3’ base pairs at positions -10 and -11 (Baek et al. 2024); the “CmR” motif (green), also known as the “GHG” motif (where ‘H’ is A, C or U) (Fang & Bartel 2015) or the “mGHG” motif (where ‘m’ is mismatch) (Kwon et al. 2019), that shows a bias for cytidine at position -5 of the 5’ DROSHA cut, a purine ---– usually a guanosine -– at position +5 of the 3’ DROSHA cut, and a base-pair mismatch between nucleotides -6 and +4 of the two DROSHA sites; the “mid-BMW” (purple) motif that reflects the propensity for bulges, wobbles and/or mismatches to occur at positions 10-12 on the mature arm relative to the star arm (Wolter et al. 2017; Li et al. 2020, 2021); the apical or distal “UGUG” motif (orange); and the “CNNC” motif (blue).

At the top of Figure 4A are the 9497 microRNA sequences annotated with a 5p mature gene product; below these are the 10940 microRNA sequences annotated with a 3p mature gene product. Co-mature microRNAs (i.e., those whose read counts for both arms that are within 2X of one another,) are not included in Figure 4 (see Supplementary Figure 2). The percentages indicate the proportion of reads for each region of the pre-microRNA including the Drosha-cleavage (DcRNA, (48), also known as microRNA offset reads (Mors, (49)), both arms, as well as the loop. In color, are the sequence propensities along the entirety of the pre-microRNA sequence, with sequence logos to highlight particularly relevant preferences in each of the two arms. As is well known, mature microRNA sequences are dramatically enriched in uridine (U) at positions one and nine, irrespective of whether they are 5p or 3p mature sequences. In contrast, the star sequences have different starting nucleotides, with 3p stars (i.e., those associated with 5p matures) typically possessing a cytosine (C) at position 1 and 5p stars typically starting with an adenosine (A). Regardless of this difference, both 3p and 5p stars underemphasize both guanosine (G) and uridine. These differences in the primary sequence propensities of 5p versus 3p matures are accompanied by differences in their secondary structure. The starting uridine in position 1 of 5p matures is generally mismatched with the corresponding nucleotide of the 3p arm, whereas the starting uridine of 3p matures is generally base paired to its cognate partner on the 5p arm (M, the darker the gray in Figure 4A the higher the mismatch rate between the specified nucleotide and its cognate on the other arm). These sequence and structural propensities, coupled with clear signals in sequence evolution (13), highlight that there are indeed important evolutionary and mechanistic distinctions not only between mature and star sequences at large, but also between mature sequences derived from the 5p arm and those derived from the 3p arm. These observations are in striking contrast to the common assumption that the arms of a microRNA locus are effectively interchangeable and, as such, not worth delineating as either mature or star products (50). This, of course, in no way means that both arm products cannot be used as regulatory molecules. Instead, it simply reflects that, on average, most microRNA loci have an evolutionarily selected and mechanistically driven mature gene product that can be discriminated from its necessary but passive partner arm based on discrete sequence and structural propensities (Figure 4A)

In addition to these characteristics of the fully processed arms, motifs in the primary and precursor microRNAs also help the microprocessor to distinguish *bona fide* pre-microRNA sequences from other transcribed (but non-microRNA) genomic hairpins and to position its two RNase III domains at the appropriate site of the pri-microRNA sequence (49, 51). Six such motifs are currently recognized, each of which disrupts the overall structural symmetry of the prototypical pri-microRNA (52) (Figure 4B), albeit with only a moderate effect on processing individually, and diminishing returns collectively (Auyeung et al. 2013; Baek et al. 2024). Two motifs help position DROSHA and DGCR8 to the ends of the double-stranded pre-microRNA: the basal “UG” motif (Figure 4B, pink), which guides DROSHA to the proximal end of the hairpin (36), and the apical “UGUG” motif (Figure 4B, orange) that guides DGCR8 to the distal end of the hairpin. The “CNNC” motif (where N = any nt), found on the single stranded end of the 3p arm (Figure 4B, blue), appears to engage the splicing factors SRSF3 and SRSF7 to assist in pre-microRNA recognition and processing (53, 54). The “mid-BWM” motif—so named because of the propensity to find bulges, wobbles and/or mismatches at positions 10-12 on the mature arm relative to the star arm (Figure 4B, purple)—also helps position DROSHA at the correct cutting site by preventing it from binding to the apical junction (55–57). The “bulged GWG” motif with a bulged uridine or adenosine (W) on the 3p arm flanked by 5’C-G-3’ base pairs at positions -10 and -11 on the 5p arm (Figure 4B, yellow) dramatically increases microprocessing efficiency and homogeneity (36). Finally, a motif located a few nucleotides upstream of the 5p DROSHA cut and a few nucleotides downstream of the 3p DROSHA cut appears to be the key positioning motif for DROSHA. This motif is called the “GHG” motif (where ‘H’ is A, C or U) by Fang & Bartel (52) or the “mGHG” motif (where ‘m’ is mismatch) by Kwon et al. (58). However, given the data in Figure 4, it appears that neither of these monikers reflects the actual sequence propensities found in this motif. Instead, the bias appears to be for a cytidine at position -5 of the 5’ DROSHA cut, a purine – usually a guanosine – at position +5 of the 3’ DROSHA cut, and a mismatch between nucleotides -6 and +4 of the two DROSHA sites. Hence, we refer to this motif as the “CmR” motif (Figure 4A, B, green) where the “C” highlights the propensity for the cytidine, the “m” the mismatch, and the “R” the purine. This motif is recognized by the dsRBD motif of DROSHA, which helps precisely position the two catalytic RNase III domains on the pri-microRNA relative to the single- to double-stranded junction (58).

With so many annotated microRNAs from such a broad collection of metazoan taxa, it is likely that any additional sequence or structural propensities whose presence can be delineated by simply comparing sequences and secondary structures among species do not exist. However, the losses of the features described above are easily discerned, most dramatically in *Caenorhabditis* nematodes. Both *C. elegans* and *C. briggsae* have lost essentially all of these motifs, replacing them with a new and unique secondary structural signal 5’ and 3’ of the ancestral CmR motif (59, 60). Because nematodes are not basal bilaterians, but instead nested within the Ecdysozoa with animals like arthropods (61), the absence of these ancestral motifs and the appearance of this new structural signal long postdates the bilaterian LCA, highlighting that motifs are part of the ancestral mode of microRNA processing that evolved long before the appearance of nematodes and, indeed, were likely present in the bilaterian LCA.

### microRNA annotation workflow and *modus operandi* of MirGeneDB

The creation of MirGeneDB was prompted by our own efforts to conduct comparative microRNA studies (40, 41, 62, 63) and the obvious lack of a well-curated microRNA reference database. For this reason, our past, current, and future selection of new taxa for MirGeneDB has been, is, and will be informed by the need to efficiently represent the broad diversity of the metazoan tree of life.

The incorporation of taxa generally requires the availability of a genome reference (either a complete assembly or raw genomic sequencing data) and smallRNA read data from the same or a very closely related species. Exceptions to this rule in MirGeneDB 3.0 primarily include evolutionarily and phylogenetically important species for which reads are currently not available and will not likely be available in the foreseeable future, such as the coelacanth, the tuatara, the chambered nautilus, and the annelid *Dimorphilus*. For these genome-only species, all reads are clearly indicated as “predicted,” and the determination of mature versus star arms was made based on the processing of the orthologous sequences in a closely related species. Assembled genomes are downloaded and preliminarily annotated by MirMachine (28) while unassembled genomic read data is processed along with smallRNA seq data by MirCandRef (24). The acquisition of publicly available smallRNA sequencing data is less straightforward because, in our experience, many existing publications either have not deposited their data in the Sequence Read Archive (SRA) (64), or suffer from poor sample annotation. This may have historical causes because data deposition was not mandatory until recently, with the SRA only being launched in 2007. These difficulties are complicated even further by the issue of orphan SRA datasets, which lack PubMed IDs because they have never been formally described in a publication, even though they are publicly available. To meet these challenges, we use miSRA (https://github.com/bioinfoUGR/miSRA), a newly developed and publicly available tool to screen the SRA repository for existing sequencing data as a function of species and other metadata of interest. Frequently, SRA metadata are scarce or even incorrect, so we manually (re)annotate the origin, accession, and publication information of all smallRNA sequencing datasets as possible. We provide all processed and annotated fasta files of smallRNA sequencing data for each species on our download section (https://master.cloud.mirgenedb.org/download) as a service to our users and the public in general (see Supplementary Table 2).

These data are either downloaded and processed with miSRA, or downloaded with SRA-toolkit (64) and processed by miRTrace (65). The processed reads, together with the MirMachine predictions of conserved microRNAs from the genome will be subject to a microRNA prediction run with MirMiner (43). The microRNA prediction results are then manually curated and annotated using synteny information (if available) and tree-based comparisons of sequence deviations. This rigorous process is time-consuming, but guarantees the highest possible quality for our microRNA annotations, which can be considered to be near-free of false positives.

To ensure that the annotation process is consistent for all species in MirGeneDB, we are not open to direct submissions of annotations, though we are always grateful for suggestions and pointers to new species and data. If we act on a suggestion for a new species, we reserve the right to re-annotate its microRNA complement before hosting it on MirGeneDB.

### Computational Improvements

The most important and noteworthy change is that as of this release MirGeneDB is hosted on two independent servers in two different cities in Norway through the Norwegian Research and Education Cloud (NREC) that serve as computational mirrors in case one of them is affected and cannot function.

### Conclusion

More than 30 years and 150, 000 publications after the discovery of microRNAs (66), the importance and relevance of these regulatory molecules for biomedical, evolutionary, and biosystematic research questions are obvious. However, determining what is and what is not a microRNA and annotating them as such has remained a computational and intellectual challenge with major uncertainties about the validity and reliability of public microRNA repositories. With the advent of next-generation sequencing, high-performance computational pipelines, and groundbreaking studies that deciphered the intricate mechanism of microRNA biogenesis (see (30, 67)), as well as their unique mode of evolution (see (25, 44)), the rules for annotation and nomenclature of microRNAs are now robustly established (13). Despite these advances, spurious annotations of thousands of questionable “microRNAs” continue to be released in an apparently never-ending race for more, rather than better data (68–71), while existing incorrect annotations have lead to a flood of highly questionable research findings (72–76), with the potential to create serious harm, both for patients and for the field as a whole, when dealing with human diseases.

Just as natural history museums provide invaluable collections for the biological research community, repositories or databases of genetic elements should provide a reliable reference to the corresponding research community. To guarantee reliability and comparability, physical natural history museums do not allow the deposition of specimens to their collections without first passing them through a rigorous curatorial process by expert taxonomists that adhere to a strict set of rules for the description (and naming) of new species (77). Repositories of genes, coding or non-coding, should also apply such rigorous and institutionalized curation. We see MirGeneDB as this institution for microRNAs.

In the decade since its 2015 release, MirGeneDB has undergone 3 major expansions that have placed it at the forefront of microRNA annotation and curation and brought it closer to the goal of complete representation of all animal phyla. All expansions adhered to the annotation and nomenclature rules initially established in 2003 (27) and later expanded to the Next Generation Sequencing level (13), introduced new species and computational features such as integration of read data or evolutionary nodes of origin for families and genes, respectively. This release, besides its many novel species, makes read data we collected from various sources over the years directly available on our website, along with updated and downloadable covariance models, as powerful tools for accurate microRNA discovery. Our new naming responsibilities will unite the curatorial and nomenclatural functions in the microRNA field and ensure that future studies of microRNA evolution and function can rely on a dynamic database that grows with the ever-increasing number of publicly available metazoan genomes and small RNA libraries.

We hope to both ignite and contribute to a renaissance in microRNA research by moving beyond an era dominated by technological advancements, which fueled a race to describe increasingly dubious microRNAs in few model systems and diseases with limited biological significance. Instead, we aim to shift the focus toward an emphasis on biology - as it was also pointed out by others (26, 34, 35) - utilizing comparative approaches based on solid curatorial foundations to deepen our understanding on the emergence of complexity of animal life and beyond.

## Supporting information

SuppFig1

SuppFig2

SuppTable2

SuppTable1

## Acknowledgement

We are grateful to Jose M Martin Duran (Queen Mary University London) for samples of *Owenia fusiformis*, Mansi Srivastava (Harvard University) for samples of Hofstenia miamia and Bernhard Egger (University of Innsbruck) for the *Prostheceraeus crozieri* specimen. We also thank Detlev Arendt (EMBL Heidelberg) for early access to the *Platynereis dumerilii* read data. We thank Blake Sweeney (EMBL-EBI, RNAcentral & Rfam) for support and help integrating MirGeneDB and Covariance models into Rfam, RNAcentral. Leanne Haggerty (EMBL-EBI, Ensembl) and Jose M. G. Perez-Silva (EMBL-EBI, Ensembl) are acknowledged for their work with MirGeneDB and MirMachine integration into Ensembl. We thank David P. Bartel (HHMI, MIT, Whitehead Institute) and Victor Ambros (UMass Chan) for discussions and encouragement on the new naming responsibilities of novel conserved microRNAs.

## Funding

AWC was funded by the Dartmouth James O. Freedman Presidential Scholars program. BF is funded by Tromsø forskningsstiftelse grant (TFS) [20_SG_BF ‘MIRevolution’].

## Data availability statement

All data herein is publicly available on SRA and on MirGeneDB as detailed in Supplementary File 1.

## Notes

### Competing Interest Statement

The authors have declared no competing interest.

## References

1. Fromm, B., Zhong, X., Tarbier, M., Friedländer, M.R. and Hackenberg, M. (2022) The limits of human microRNA annotation have been met. RNA, 28, 781–785.

2. Fromm, B., Patil, A.H. and Halushka, M.K. (2022) A Novel Circulating MicroRNA for the Detection of Acute Myocarditis. N. Engl. J. Med., 387, 1240.

3. McIlwraith, E.K., He, W. and Belsham, D.D. (2023) Promise and perils of MicroRNA discovery research: Working toward quality over quantity. Endocrinology, 164.

4. Castellano, L. and Stebbing, J. (2013) Deep sequencing of small RNAs identifies canonical and non-canonical miRNA and endogenous siRNAs in mammalian somatic tissues. Nucleic Acids Res., 41, 3339–3351.

5. Chiang, H.R., Schoenfeld, L.W., Ruby, J.G., Auyeung, V.C., Spies, N., Baek, D., Johnston, W.K., Russ, C., Luo, S., Babiarz, J.E., et al. (2010) Mammalian microRNAs: experimental evaluation of novel and previously annotated genes. Genes Dev., 24, 992–1009.

6. Jones-Rhoades, M.W. (2012) Conservation and divergence in plant microRNAs. Plant Mol. Biol., 80, 3–16.

7. Ludwig, N., Becker, M., Schumann, T., Speer, T., Fehlmann, T., Keller, A. and Meese, E. (2017) Bias in recent miRBase annotations potentially associated with RNA quality issues. Sci. Rep., 7, 5162.

8. Langenberger, D., Bartschat, S., Hertel, J., Hoffmann, S., Tafer, H. and Stadler, P.F. (2011) MicroRNA or Not MicroRNA? In Advances in Bioinformatics and Computational Biology. Springer Berlin Heidelberg, pp. 1–9.

9. Meng, Y., Shao, C., Wang, H. and Chen, M. (2012) Are all the miRBase-registered microRNAs true? A structure- and expression-based re-examination in plants. RNA Biol., 9, 249– 253.

10. Tarver, J.E., Donoghue, P.C. and Peterson, K.J. (2012) Do miRNAs have a deep evolutionary history? Bioessays, 34, 857–866.

11. Taylor, R.S., Tarver, J.E., Hiscock, S.J. and Donoghue, P.C. (2014) Evolutionary history of plant microRNAs. Trends Plant Sci., 10.1016/j.tplants.2013.11.008.

12. Wang, X. and Liu, X.S. (2011) Systematic Curation of miRBase Annotation Using Integrated Small RNA High-Throughput Sequencing Data for C. elegans and Drosophila. Front. Genet., 2, 25.

13. Fromm, B., Billipp, T., Peck, L.E., Johansen, M., Tarver, J.E., King, B.L., Newcomb, J.M., Sempere, L.F., Flatmark, K., Hovig, E., et al. (2015) A Uniform System for the Annotation of Vertebrate microRNA Genes and the Evolution of the Human microRNAome.

14. Axtell, M.J. and Meyers, B.C. (2018) Revisiting Criteria for Plant MicroRNA Annotation in the Era of Big Data. Plant Cell, 30, 272–284.

15. Guo, Z., Kuang, Z., Wang, Y., Zhao, Y., Tao, Y., Cheng, C., Yang, J., Lu, X., Hao, C., Wang, T., et al. (2020) PmiREN: a comprehensive encyclopedia of plant miRNAs. Nucleic Acids Res., 48, D1114–D1121.

16. Fromm, B., Keller, A., Yang, X., Friedlander, M.R., Peterson, K.J. and Griffiths-Jones, S. (2020) Quo vadis microRNAs? Trends Genet., 36, 461–463.

17. Fromm, B., Høye, E., Domanska, D., Zhong, X., Aparicio-Puerta, E., Ovchinnikov, V., Umu, S.U., Chabot, P.J., Kang, W., Aslanzadeh, M., et al. (2022) MirGeneDB 2.1: toward a complete sampling of all major animal phyla. Nucleic Acids Res., 50, D204–D210.

18. Fromm, B., Domanska, D., Høye, E., Ovchinnikov, V., Kang, W., Aparicio-Puerta, E., Johansen, M., Flatmark, K., Mathelier, A., Hovig, E., et al. (2020) MirGeneDB 2.0: the metazoan microRNA complement. Nucleic Acids Res., 48, D132–D141.

19. Budak, H., Bulut, R., Kantar, M. and Alptekin, B. (2016) MicroRNA nomenclature and the need for a revised naming prescription. Brief. Funct. Genomics, 15, 65–71.

20. Ruby, J.G., Jan, C., Player, C., Axtell, M.J., Lee, W., Nusbaum, C., Ge, H. and Bartel, D.P. (2006) Large-scale sequencing reveals 21U-RNAs and additional microRNAs and endogenous siRNAs in C. elegans. Cell, 127, 1193–1207.

21. Ruby, J.G., Stark, A., Johnston, W.K., Kellis, M., Bartel, D.P. and Lai, E.C. (2007) Evolution, biogenesis, expression, and target predictions of a substantially expanded set of Drosophila microRNAs. Genome Res., 17, 1850–1864.

22. Grimson, A., Srivastava, M., Fahey, B., Woodcroft, B.J., Chiang, H.R., King, N., Degnan, B.M., Rokhsar, D.S. and Bartel, D.P. (2008) Early origins and evolution of microRNAs and Piwi-interacting RNAs in animals. Nature, 455, 1193–1197.

23. Jan, C.H., Friedman, R.C., Ruby, J.G. and Bartel, D.P. (2011) Formation, regulation and evolution of Caenorhabditis elegans 3’UTRs. Nature, 469, 97–101.

24. Fromm, B., Worren, M.M., Hahn, C., Hovig, E. and Bachmann, L. (2013) Substantial loss of conserved and gain of novel MicroRNA families in flatworms. Mol. Biol. Evol., 30, 2619– 2628.

25. Fromm, B. (2024) A renaissance of microRNAs as taxonomic and phylogenetic markers in animals. Zool. Scr., 10.1111/zsc.12684.

26. Witwer, K.W. and Halushka, M.K. (2016) Toward the promise of microRNAs - Enhancing reproducibility and rigor in microRNA research. RNA Biol., 13, 1103–1116.

27. Ambros, V., Bartel, B., Bartel, D.P., Burge, C.B., Carrington, J.C., Chen, X., Dreyfuss, G., Eddy, S.R., Griffiths-Jones, S., Marshall, M., et al. (2003) A uniform system for microRNA annotation. RNA, 9, 277–279.

28. Umu, S.U., Paynter, V.M., Trondsen, H., Buschmann, T., Rounge, T.B., Peterson, K.J. and Fromm, B. (2023) Accurate microRNA annotation of animal genomes using trained covariance models of curated microRNA complements in MirMachine. Cell Genom, 3, 100348.

29. Langschied, F., Leisegang, M.S., Brandes, R.P. and Ebersberger, I. (2023) ncOrtho: efficient and reliable identification of miRNA orthologs. Nucleic Acids Res., 51, e71.

30. Bartel, D.P. (2018) Metazoan MicroRNAs. Cell, 173, 20–51.

31. Kang, W., Fromm, B., Houben, A.J., Høye, E., Bezdan, D., Arnan, C., Thrane, K., Asp, M., Johnson, R., Biryukova, I., et al. (2021) MapToCleave: High-throughput profiling of microRNA biogenesis in living cells. Cell Rep., 37, 110015.

32. Kim, K., Baek, S.C., Lee, Y.-Y., Bastiaanssen, C., Kim, J., Kim, H. and Kim, V.N. (2021) A quantitative map of human primary microRNA processing sites. Mol. Cell, 81, 3422–3439.e11.

33. Patil, A.H., Baran, A., Brehm, Z.P., McCall, M.N. and Halushka, M.K. (2022) A curated human cellular microRNAome based on 196 primary cell types. Gigascience, 11.

34. Johnson, K.C., Johnson, S.T., Liu, J., Chu, Y. and Corey, D.R. (2022) Prioritizing annotated miRNAs: Only a small percentage are candidates for biological regulation. 10.1101/2022.10.18.512653.

35. Johnson, K.C., Johnson, S.T., Liu, J., Chu, Y., Arana, C., Han, Y., Wang, T. and Corey, D.R. (2023) Consequences of depleting TNRC6, AGO, and DROSHA proteins on expression of microRNAs. RNA, 29, 1166–1184.

36. Baek, S.C., Kim, B., Jang, H., Kim, K., Park, I.-S., Min, D.-H. and Kim, V.N. (2024) Structural atlas of human primary microRNAs generated by SHAPE-MaP. Mol. Cell, 10.1016/j.molcel.2024.02.005.

37. Le, C.T., Nguyen, T.D. and Nguyen, T.A. (2024) Two-motif model illuminates DICER cleavage preferences. Nucleic Acids Res., 52, 1860–1877.

38. Hagemann, L., Mauer, K.M., Hankeln, T., Schmidt, H. and Herlyn, H. (2023) Nuclear genome annotation of wheel animals and thorny-headed worms: inferences about the last common ancestor of Syndermata (Rotifera s.l.). Hidrobiol. (Sofiâ), 10.1007/s10750-023-05268-6.

39. Poulin, R. and Randhawa, H.S. (2015) Evolution of parasitism along convergent lines: from ecology to genomics. Parasitology, 142 **Suppl 1**, S6–S15.

40. Herlyn, H., Hembrom, A.A., Tosar, J., Mauer, K.M., Schmidt, H., Dezfuli, B.S., Hankeln, T., Bachmann, L., Sarkies, P., Peterson, K.J., et al. (2024) Substantial hierarchical reductions of genetic and morphological traits in the evolution of rotiferan parasites. bioRxiv, 10.1101/2024.08.01.605096.

41. Peterson, K.J., Clarke, A., Zolotarov, G., Deline, B., McPeek, M., Martinez, P. and Fromm, B. (2024) Capturing changes to animal complexity from quantifiable patterns in genomic data. bioRxiv, 10.1101/2024.08.22.609214.

42. Sempere, L.F., Cole, C.N., McPeek, M.A. and Peterson, K.J. (2006) The phylogenetic distribution of metazoan microRNAs: insights into evolutionary complexity and constraint. J. Exp. Zool. B Mol. Dev. Evol., 306, 575–588.

43. Wheeler, B.M., Heimberg, A.M., Moy, V.N., Sperling, E.A., Holstein, T.W., Heber, S. and Peterson, K.J. (2009) The deep evolution of metazoan microRNAs. Evol. Dev., 11, 50– 68.

44. Tarver, J.E., Sperling, E.A., Nailor, A., Heimberg, A.M., Robinson, J.M., King, B.L., Pisani, D., Donoghue, P.C.J. and Peterson, K.J. (2013) miRNAs: small genes with big potential in metazoan phylogenetics. Mol. Biol. Evol., 30, 2369–2382.

45. Seitz, H., Royo, H., Bortolin, M.-L., Lin, S.-P., Ferguson-Smith, A.C. and Cavaillé, J. (2004) A large imprinted microRNA gene cluster at the mouse Dlk1-Gtl2 domain. Genome Res., 14, 1741–1748.

46. Kozomara, A., Birgaoanu, M. and Griffiths-Jones, S. (2019) miRBase: from microRNA sequences to function. Nucleic Acids Res., 47, D155–D162.

47. Gao, Z., Wang, M., Blair, D., Zheng, Y. and Dou, Y. (2014) Phylogenetic analysis of the endoribonuclease Dicer family. PLoS One, 9, e95350.

48. Ma, H., Wu, Y., Choi, J.-G. and Wu, H. (2013) Lower and upper stem-single-stranded RNA junctions together determine the Drosha cleavage site. Proc. Natl. Acad. Sci. U. S. A., 110, 20687–20692.

49. Shi, W., Hendrix, D., Levine, M. and Haley, B. (2009) A distinct class of small RNAs arises from pre-miRNA-proximal regions in a simple chordate. Nat. Struct. Mol. Biol., 16, 183– 189.

50. Kozomara, A. and Griffiths-Jones, S. (2014) miRBase: annotating high confidence microRNAs using deep sequencing data. Nucleic Acids Res., 42, D68–73.

51. Gu, K., Mok, L., Wakefield, M.J. and Chong, M.M.W. (2024) Non-canonical RNA substrates of Drosha lack many of the conserved features found in primary microRNA stem-loops. Sci. Rep., 14, 6713.

52. Fang, W. and Bartel, D.P. (2015) The Menu of Features that Define Primary MicroRNAs and Enable De Novo Design of MicroRNA Genes. Mol. Cell, 60, 131–145.

53. Kim, K., Nguyen, T.D., Li, S. and Nguyen, T.A. (2018) SRSF3 recruits DROSHA to the basal junction of primary microRNAs. RNA, 24, 892–898.

54. Le, M.N., Nguyen, T.D. and Nguyen, T.A. (2023) SRSF7 and SRSF3 depend on RNA sequencing motifs and secondary structures to regulate Microprocessor. Life Sci. Alliance, 6.

55. Wolter, J.M., Le, H.H.T., Linse, A., Godlove, V.A., Nguyen, T.-D., Kotagama, K., Lynch, A., Rawls, A. and Mangone, M. (2017) Evolutionary patterns of metazoan microRNAs reveal targeting principles in the let-7 and miR-10 families. Genome Res., 27, 53–63.

56. Li, S., Nguyen, T.D., Nguyen, T.L. and Nguyen, T.A. (2020) Mismatched and wobble base pairs govern primary microRNA processing by human Microprocessor. Nat. Commun., 11, 1926.

57. Li, S., Le, T.N.-Y., Nguyen, T.D., Trinh, T.A. and Nguyen, T.A. (2021) Bulges control pri-miRNA processing in a position and strand-dependent manner. RNA Biol., 18, 1716–1726.

58. Kwon, S.C., Baek, S.C., Choi, Y.-G., Yang, J., Lee, Y.-S., Woo, J.-S. and Kim, V.N. (2019) Molecular Basis for the Single-Nucleotide Precision of Primary microRNA Processing. Mol. Cell, 73, 505–518.e5.

59. Nguyen, T.L., Nguyen, T.D., Ngo, M.K., Le, T.N.-Y. and Nguyen, T.A. (2023) Noncanonical processing by animal Microprocessor. Mol. Cell, 83, 1810–1826.e8.

60. Nguyen, T.L., Nguyen, T.D., Ngo, M.K. and Nguyen, T.A. (2023) Dissection of the Caenorhabditis elegans Microprocessor. Nucleic Acids Res., 51, 1512–1527.

61. Aguinaldo, A.M., Turbeville, J.M., Linford, L.S., Rivera, M.C., Garey, J.R., Raff, R.A. and Lake, J.A. (1997) Evidence for a clade of nematodes, arthropods and other moulting animals. Nature, 387, 489–493.

62. Zolotarov, G., Fromm, B., Legnini, I., Ayoub, S., Polese, G., Maselli, V., Chabot, P.J., Vinther, J., Styfhals, R., Seuntjens, E., et al. (2022) MicroRNAs are deeply linked to the emergence of the complex octopus brain. Sci Adv, 8, eadd9938.

63. Peterson, K.J., Beavan, A., Chabot, P., McPeek, M.L., Pisani, D., Fromm, B. and Simakov, O. microRNAs as Indicators into the Causes and Consequences of Whole Genome Duplication Events. 10.1101/2021.09.01.458616.

64. Leinonen, R., Sugawara, H., Shumway, M. and International Nucleotide Sequence Database Collaboration (2011) The sequence read archive. Nucleic Acids Res., 39, D19–21.

65. Kang, W., Eldfjell, Y., Fromm, B., Estivill, X., Biryukova, I. and Friedländer, M.R. (2018) miRTrace reveals the organismal origins of microRNA sequencing data. Genome Biol., 19, 213.

66. Lee, R.C., Feinbaum, R.L. and Ambros, V. (1993) The C. elegans heterochronic gene lin-4 encodes small RNAs with antisense complementarity to lin-14. Cell, 75, 843–854.

67. Shang, R., Lee, S., Senavirathne, G. and Lai, E.C. (2023) microRNAs in action: biogenesis, function and regulation. Nat. Rev. Genet., 24, 816–833.

68. Jha, A., Panzade, G., Pandey, R. and Shankar, R. (2015) A legion of potential regulatory sRNAs exists beyond the typical microRNAs microcosm. Nucleic Acids Res., 43, 8713– 8724.

69. Londin, E., Loher, P., Telonis, A.G., Quann, K., Clark, P., Jing, Y., Hatzimichael, E., Kirino, Y., Honda, S., Lally, M., et al. (2015) Analysis of 13 cell types reveals evidence for the expression of numerous novel primate- and tissue-specific microRNAs. Proc. Natl. Acad. Sci. U. S. A., 112, E1106–15.

70. Alles, J., Fehlmann, T., Fischer, U., Backes, C., Galata, V., Minet, M., Hart, M., Abu-Halima, M., Grässer, F.A., Lenhof, H.-P., et al. (2019) An estimate of the total number of true human miRNAs. Nucleic Acids Res., 47, 3353–3364.

71. Lorenzi, L., Chiu, H.-S., Avila Cobos, F., Gross, S., Volders, P.-J., Cannoodt, R., Nuytens, J., Vanderheyden, K., Anckaert, J., Lefever, S., et al. (2021) The RNA Atlas expands the catalog of human non-coding RNAs. Nat. Biotechnol., 39, 1453–1465.

72. Blanco-Domínguez, R., Sánchez-Díaz, R., de la Fuente, H., Jiménez-Borreguero, L.J., Matesanz-Marín, A., Relaño, M., Jiménez-Alejandre, R., Linillos-Pradillo, B., Tsilingiri, K., Martín-Mariscal, M.L., et al. (2021) A Novel Circulating MicroRNA for the Detection of Acute Myocarditis. N. Engl. J. Med., 384, 2014–2027.

73. Chinnappa, K., Cárdenas, A., Prieto-Colomina, A., Villalba, A., Márquez-Galera, Á., Soler, R., Nomura, Y., Llorens, E., Tomasello, U., López-Atalaya, J.P., et al. (2022) Secondary loss of miR-3607 reduced cortical progenitor amplification during rodent evolution. Sci Adv, 8, eabj4010.

74. Garcia-Martin, R., Wang, G., Brandão, B.B., Zanotto, T.M., Shah, S., Kumar Patel, S., Schilling, B. and Kahn, C.R. (2022) MicroRNA sequence codes for small extracellular vesicle release and cellular retention. Nature, 601, 446–451.

75. Ying, W., Gao, H., Dos Reis, F.C.G., Bandyopadhyay, G., Ofrecio, J.M., Luo, Z., Ji, Y., Jin, Z., Ly, C. and Olefsky, J.M. (2021) MiR-690, an exosomal-derived miRNA from M2-polarized macrophages, improves insulin sensitivity in obese mice. Cell Metab., 33, 781–790.e5.

76. Rohm, T.V., Castellani Gomes Dos Reis, F., Isaac, R., Murphy, C., Cunha E, Rocha, K., Bandyopadhyay, G., Gao, H., Libster, A.M., Zapata, R.C., Lee, Y.S., et al. (2024) Author Correction: Adipose tissue macrophages secrete small extracellular vesicles that mediate rosiglitazone-induced insulin sensitization. Nat. Metab., 6, 1646.

77. The ICZN amendment of articles 8, 9, 10, 21 and 78 of the International Code of Zoological Nomenclature (2012) Zoosyst. Ross., 21, 323–327.

