## Supplementary figures and images for "MirGeneDB 3.0: Improved taxonomic sampling, uniform nomenclature of novel conserved microRNA families, and updated covariance models"

### SuppFig1

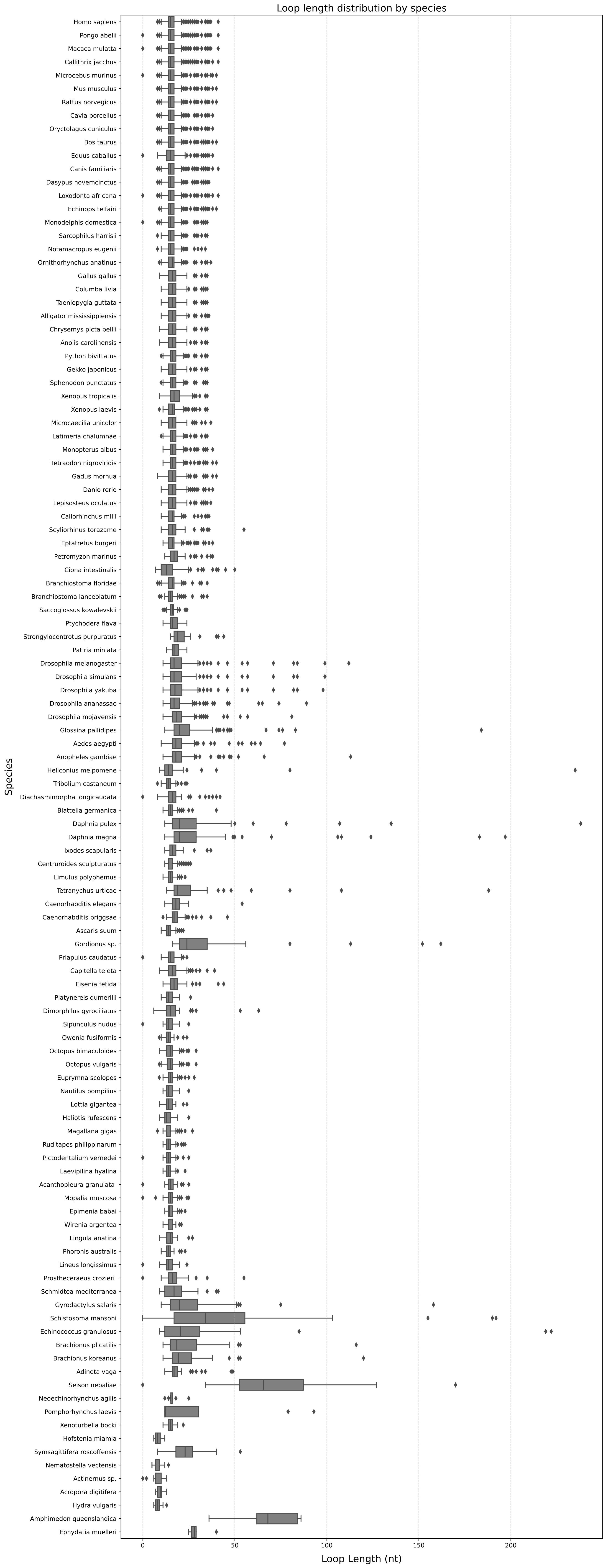

### SuppFig2

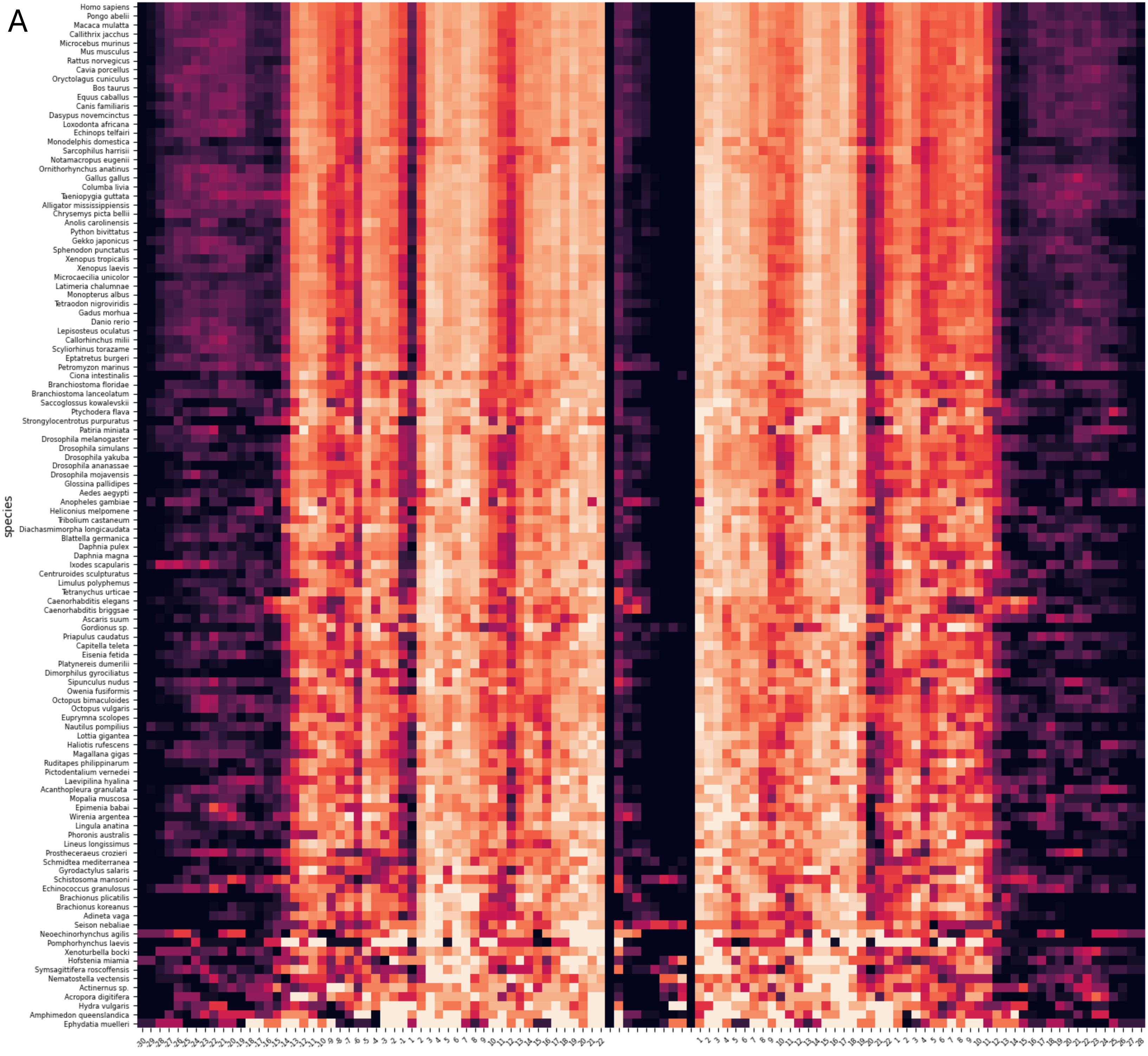

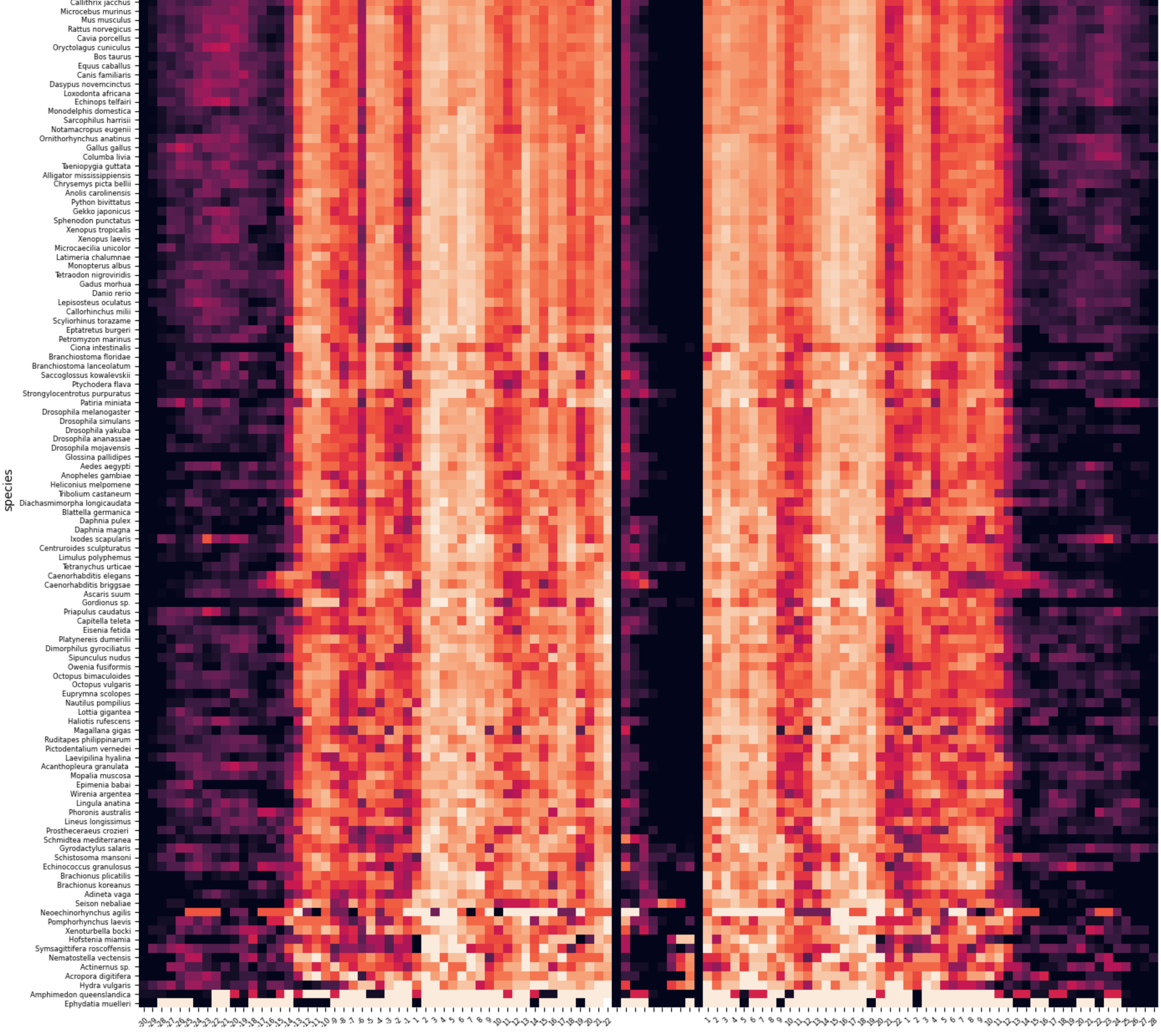

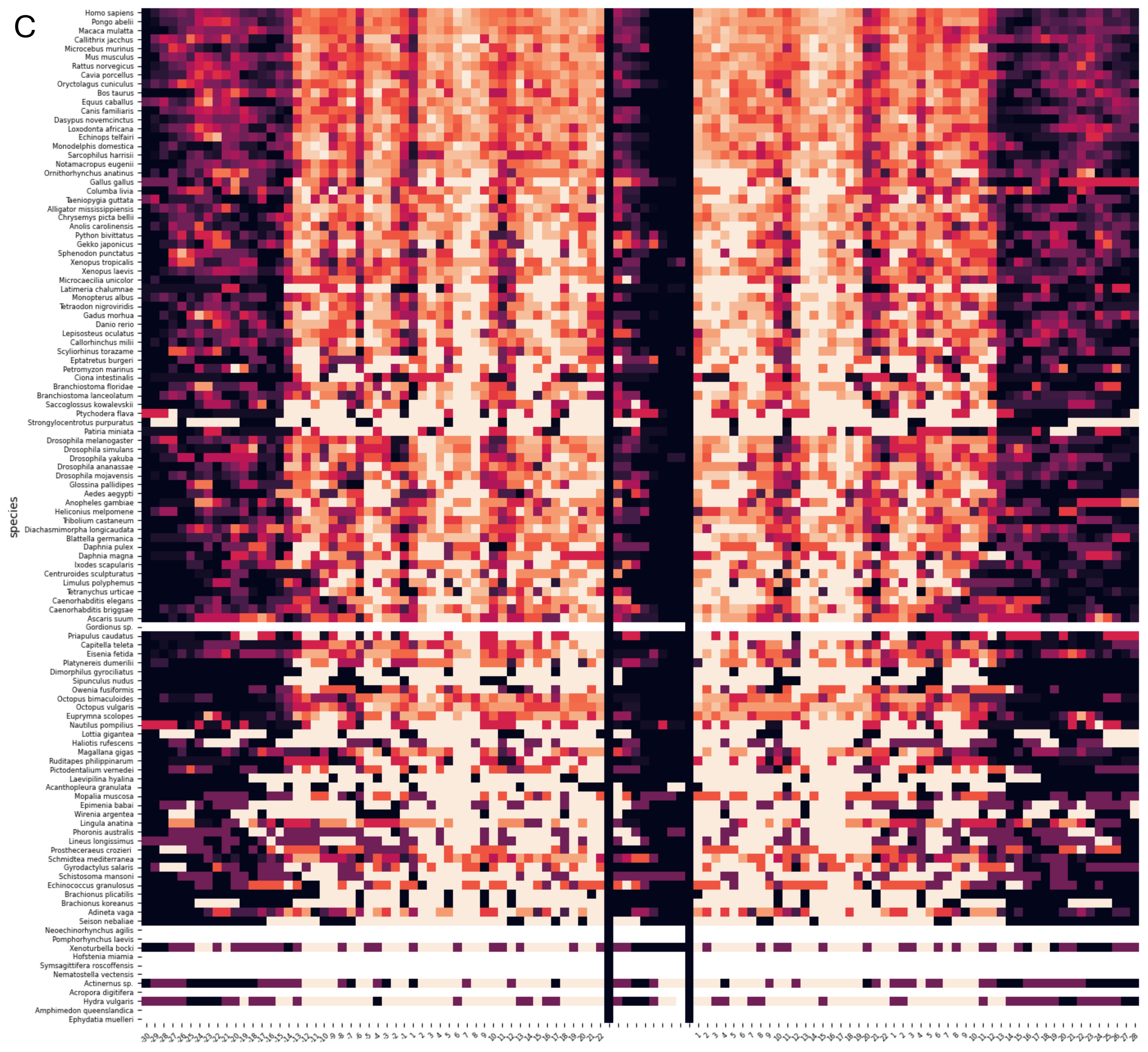
